# Immune responses in mice induced by multi-epitope DNA vaccine and protein vaccine of Crimean-Congo Hemorrhagic Fever Virus

**DOI:** 10.1101/719724

**Authors:** Meilipaiti Yusufu, Alai Shalitanati, Huan Yu, Abulimiti Moming, Yijie Li, Fei Deng, Yujiang Zhang, Surong Sun

## Abstract

Crimean-Congo Hemorrhagic Fever (CCHF), caused by the CCHF virus (CCHFV), is a severe tick borne zoonosis widely distributed in over 30 countries and regions. Currently, there is no licensed vaccine available for CCHF in China. To evaluate the cellular and humoral immune responses induced by multi-epitope DNA and protein vaccine of CCHF in BALB/c mice, a multi-epitope gene (MEPX) segment with tandem including six highly conservative and immunedominant B cell epitopes was designed based on the analysis of hydrophilicity and antigenic determinant sites in amino acid sequences of nucleoprotein and glycoprotein from CCHFV strain YL04057. The single and double-copy multi-epitope gene (MEPX and MEPX_2_) were respectively cloned into the eukaryotic expression vector pVAX I to construct the recombinant (r) plasmid pVAX-MEPX and pVAX-MEPX_2_ as DNA vaccines. The results of immunofluorescence in vitro showed that the pVAX-MEPX and pVAX-MEPX_2_ could be expressed in 293T cells. The recombinant prokaryotic plasmid pET-32a-MEPX and pET-32a-MEPX_2_ constructed previously were transformed them into *E. co*li BL21 (DE3), and recombinant multi-epitope proteins (rMEPX and rMEPX_2_) were obtained and purificated by Nickel affinity chromatography. Western blot results showed that rMEPX and rMEPX_2_ had good antigenicity. BALB/c mice were immunized with DNA vaccine alone, protein vaccine alone, and DNA prime followed by recombinant protein boost immunization strategy, respectively. After three immunizations, MTT assay, cytokine content assay, and ELISA assay for antibody titers were used to evaluate the immune response. The proliferation of mouse specific T lymphocytes in the enhanced by pVAX-MEPX_2_ combined with rMEPX_2_ boosting group was significant, and the expression levels of serum IFN-γ and IL-4 in mice were as high as 118.67 pg/mL and 135.33 pg/mL with significant difference compared to the control group (*p*<0.01), and serum antibody titer could reach up to 4.1×10^5^. Double-copy multi-epitope vaccines groups (pVAX-MEPX_2_+ rMEPX_2_) generated better cellular and humoral immune responses by DNA prime-protein vaccine boost combinatorial immunization. This result could lay the foundation for the development of CCHFV multi-epitope vaccine candidates.

## Introduction

Crimean-Congo Hemorrhagic Fever (CCHF) is an acute tick-borne zoonosis, which is caused by the Crimean-Congo Hemorrhagic Fever Virus (CCHFV). This virus is a member of the genus *Nairovirus* in the family Bunyaviridae [1]. It has been reported that 5 - 40% of patients show severe bleeding, resulting in shock and even death [1]. In China, the first case of CCHF was reported in 1965. Thus, this disease is also known as Xinjiang Hemorrhagic Fever (XHF). At present, the virus strains have been isolated from patients in the Bachu region of Xinjiang, where it is considered to be the region with the highest incidence of CCHF in China [1-3]. Because of the high virulence and high pathogenicity of CCHFV, this research needs to be conducted in the high-level laboratories. As a result, the suitable animal model was lacked, which limits the in-depth study of effective diagnostic, prevention, and therapeutic methods for CCHF [4]. So far, there is no licensed vaccine available for CCHF in China [5].

New generation vaccines, such as recombinant subunit proteins and DNA vaccines, are gradually replacing the traditional vaccines [6, 7]. For multi-epitope-based vaccines, the conserved immunedominant and neutralizing epitopes are typically selected for tandem expression, thereby reducing the interference and inhibition of extraneous sequences [8, 9]. Because of its low cost, easy operation, high safety, and highly effective protection, it has attracted wide attention [9]. It has also been reported that the vaccine that is made by a multi-epitope concatenated tandem of multiple epitope constituent is more effective than a single epitope vaccine [10, 11]. Therefore, the related study on multi-epitope gene tandem and its expression activity is of great significance for the development of novel epitope vaccines for viruses.

Our group previously investigated the antigenicity of the double-copy Multi-Epitope Peptide (MEPX_2_) formed in tandem by 6 linear B-cell epitopes on the CCHFV nucleoprotein NP and glycoproteins GP (Gn and Gc) [12,13]. In order to further understand whether the multi-epitope gene is immunogenic, in this work, we constructed a multi-epitope gene eukaryotic expression plasmid and identified its expression in vitro. We also used DNA vaccine alone, protein vaccine alone, and DNA prime followed by recombinant protein boost immunization strategy to immunize the BALB/c mice. Their immune response was evaluated. Our results could provide the basis to develop a novel multi-epitope vaccine for CCHFV in the future.

## Materials and Method

### Ethics statement

The study was approved by the Committee on the Ethics of Animal Experiments of Xinjiang Key Laboratory of Biological Resources and Genetic Engineering (BRGE-AE001), Xinjiang University.

### Analysis of the structure and antigen epitope of MEPX_2_

SignalP4.1 and TMHHMM online analysis software were used to analyze signal peptides and transmembrane regions of the amino acid sequence of MEPX_2_. SignalP 4.1 server predicts the presence and location of signal peptide cleavage sites in amino acid sequences from different organisms. The method incorporates a prediction of cleavage sites and a signal peptide/non-signal peptide prediction based on a combination of several artificial neural networks. DNAStar-Protean software was used to predict the possible BCEs on MEPX_2_. The prediction of secondary structure was based on Garnier and Robson[14], Chou and Fasman.[15] The hydrophilic scheme, flexible regimen, surface accessibility regimen and antigenicity index were analyzed and predicted using methods of the Kyte-Doolittle[16], Karplus-Schulz[17], Emini and Jameson-Wolf[18], respectively. Based on the results obtained from these analyses, peptides with good hydrophilicity, high accessibility, high flexibility and strong antigenicity were selected as epitope candidates.

### Mice and cell line

Female 6-8 week-old BALB/c mice were purchased from the Weitonglihua Animal Technology Co. Ltd. (Beijing, China). Human embryonic kidneys (HEK) 293T cells were stored by this research group.

### Plasmids and antibodies

pVAX-Gc-NP2, pET-32a-MEPX, pET-32a-MEPX_2_ previously constructed were stored by this research group [13, 19]. Both healthy and CCHFV-infected sheep serum samples were provided by Professor Yujiang Zhang of Xinjiang Centers for Disease Control and Prevention. Serum samples of sheep infected with CCHFV were previously identified using indirect immunofluorescent assays (IFAs) and reverse transcription polymerase chain reactions (RT-PCRs) [20].

### Other reagents and materials

RPMI-1640 culture medium was purchased from Thermo Fisher Scientific (Massachusetts, USA). Lipofectamine 2000 was purchased from Invitrogen (California, USA). Complete and incomplete Freund’s adjuvant and MTT (Methyl Thiazolyl Tetrazolium) were purchased from Sigma (San Francisco, USA). Mice IL-4 and IFN-γ ELISA kit was purchased from Wuhan BOSTER Biological Technology Co. Ltd. High-purity endotoxin-free plasmid extraction kit was purchased from Biotech Engineering Co., Ltd (Shanghai, China). Enhanced chemiluminescence (ECL) plus western blotting detection kit (GE Healthcare, Buckinghamshire, UK) were obtained.

### Construction of eukaryotic expression plasmids

In our previous study, a multi-epitope gene (MEPX) segment with tandem including six highly conservative and dominant B cell epitopes (NP^245-259^, NP^265-273^, Gn^39-58^, Gn^119-154^, Gc^226-245^, Gc^503-530^) was designed based on the analysis of hydrophilicity and antigenic determinant sites in amino acid sequences of nucleoprotein and glycoprotein from CCHFV strain YL04057 [13]. The epitopes in MEPX were separated from each other by a flexible peptide Gly-Gly [13]. Based on the single-copy poly-epitope gene sequence designed by Yu Huan in our group [13], we have designed a double-copy multi-epitope gene sequence optimized by E. coli preference codon (Fig 1(A)). The single-copy gene sequence MEPX and the double-copy gene sequence MEPX_2_ was synthesized by Jierui Bioengineering Co. Ltd. in Shanghai, and inserted into the expression plasmid pVAXI by restriction endonuclease digestion with *Bam*HI and *Xho*I, respectively. After being transformed into competent *Escherichia coli* DH5α cells, the recombinant plasmid pVAX-MEPX, pVAX-MEPX_2_ was identified using restricted enzyme analysis and sequencing (Sangon Biotech, Shanghai, China).

**Fig 1.**
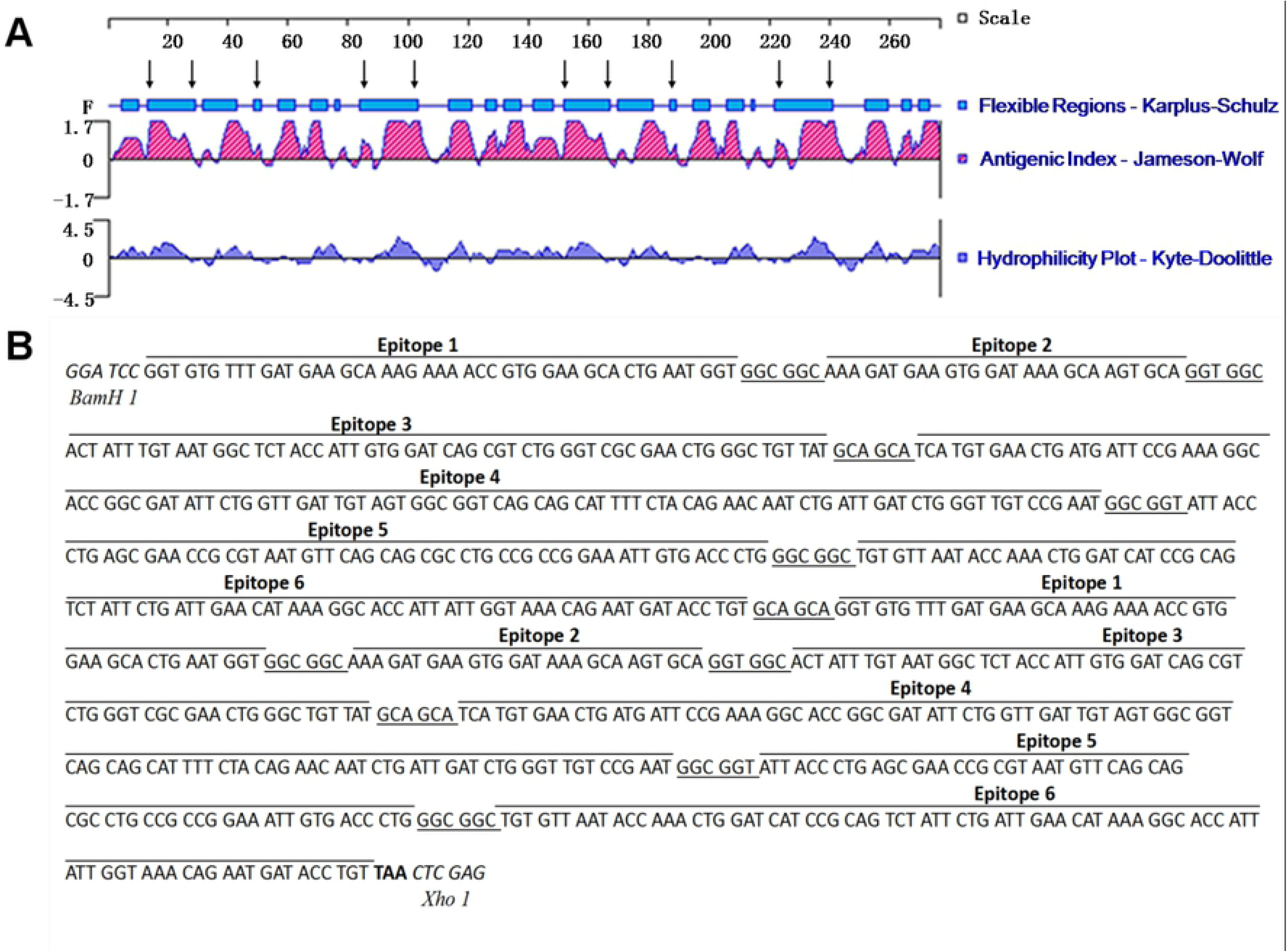
The nucleotide sequence and the bioinformatics analysis of multi-epitope gene MEPX_2_ of CCHFV. (A) The nucleotide sequence of multi-epitope gene MEPX_2_ of CCHFV. The nucleotide sequence in italicized bold is the restriction site, the nucleotide sequence within the border is the stop codon, the underlined nucleotide sequence is a flexible amino acid that links each epitope, and the remaining nucleotide sequence is the nucleotide sequence encoding the dominant epitope. (B) The secondary structure of multi-epitope gene MEPX_2_ of Gn sequence of the CCHFV YL040057 strain using DNAStar Protean software.

### Transfection and identification of recombination plasmid

HEK 293T cells cultured in 24-well plats were transiently transfected with 40 μg of the endo-toxin-free plasmids pVAX-MEPX, pVAX-MEPX_2_, pVAXI (negative control) and pVAX-Gc-NP2 (positive control) [19] using Lipofectamine 2000 (Invitrogen, USA). The cells were collected 24 h post-transfection for analysis of protein expression by indirect immunofluorescence (IFA). Briefly, the cells were fixed and permeabilized, the mixed anti-Gn, Gc, NP rabbit sera (1:100) were used as the primary antibody, and FITC-conjugated goat anti-rabbit IgG (1:2000) (TransGen Biotech, Beijing, China) as used was the secondary antibody, finally the effects were visualized under a fluorescent microscope (Leica, Germany).

### Purification and identification of the recombination protein

The overnight cultures of pET-32a-MEPX/BL21 and pET-32a-MEPX_2_/BL21 in *Escherichia coli* BL21cells were induced by IPTG and expressed at optimal expression levels and temperature. The bacteria were collected and suspended in PBS, followed by sonication at 4 °C. To optimize the purification condition using Nickel affinity chromatography, the expressed recombinant proteins (rMEPX and rMEPX_2_) from the inclusion bodies were added to a Ni affinity chromatography column and reacted for 3 h at 4°C. After the column was washed sequentially with increasing concentrations of imidazole for optimization, the concentration of 120 mmol/L imidazole was determined to be optimal for washing the target protein. The concentrations of protein were measured using bicinchonininc acid (BCA) kit (Thermo scientific). The identity of the expressed and purified recombinant protein was analyzed by SDS-PAGE and Western blot. A mouse anti-His monoclonal antibody (1: 2000), CCHFV-positive sheep serum (1:100) was used as the primary antibody, and horseradish peroxidase (HRP)-conjugated goat anti-mouse/ mouse anti-goat IgG (TransGen Biotech, Beijing, China) was used as the secondary antibody. The bound antibodies were observed through incubating membranes using ECL chmilumiescence reagents, and then were exposed in the LAS 4000 Ultrasensitive Chemiluminescence.

### Mouse immunization

Mice were randomly divided into 8 groups (Table 1), which were immunized on day 0, 14, and 28. Mice in pVAX-MEPX and pVAX-MEPX_2_ groups were respectively immunized intramuscularly (i.m) with DNA vaccine in mouse legs using an electric pulse gene transfection device. Mice in group pVAXI were immunized with empty plasmid pVAXI as a negative control. All of the DNA vaccines immunization groups were immunized with 100 μg plasmid (diluted in 100 μL PBS) with an equivalent dose three times at biweekly interval.

**Table 1.** BALB/c mouse immunization strategy. Overview of mice immunization groups.

| Group | No. of mices | Immune mode | Immune interval |
| --- | --- | --- | --- |
| pVAX-MEPX | 8 | Intramuscular<br>injection | 14 d |
| pVAX-MEPX<sub>2</sub> | 8 | Intramuscular<br>injection | 14 d |
| pVAX I | 8 | Intramuscular<br>injection | 14 d |
| rMEPX | 8 | Subcutaneous<br>injection | 14 d |
| rMEPX<sub>2</sub> | 8 | Subcutaneous<br>injection | 14 d |
| PBS | 8 | Subcutaneous<br>injection | 14 d |
| pVAX-MEPX+rMEPX | 8 | Intramuscular<br>injection<br>+<br>Subcutaneous<br>injection | 14 d |
| pVAX-MEPX<sub>2</sub>+rMEPX<sub>2</sub> | 8 | Intramuscular<br>injection<br>+<br>Subcutaneous<br>injection | 14 d |

Mice in groups rMEPX and rMEPX_2_ were respectively immunized with the purified recombinant protein. The back of the mice injected subcutaneously into 300 μg recombinant protein diluted in 300 μL PBS with an equal volume of Freund’s adjuvant. Freund’s complete adjuvant FCA was used for primary immunization and Freund’s incomplete adjuvant FIA was used for booster immunization. Group PBS were immunized i.m with an equal volume of filter-sterilized PBS as negative control group. In the group pVAX-MEPX+rMEPX, pVAX-MEPX_2_+rMEPX_2_ as the DNA prime-protein vaccine boost combinatorial immunization was administered: the primary immunization with plasmid DNA was performed, and then 14 d and 28 d after the first immunization, the recombinant protein were immunized i.m as the boosting immunization, respectively. The serum samples were collected by the mice eyelids bleeding before the each administration and 14 d after the last immunization, followed by storage at −20 °C for antibody detection by ELISA. Splenocytes were harvested for splenocyte proliferation and cytokine assay on 14 d after final immunization. The specific immunization groups and strategy was shown in the table 1 below.

### ELISA

Antigen-specific IgG antibodies (Abs) titers were measured by ELISA. Serum samples were diluted at 1:100 in the blocking agent, respectively. Briefly, 96-well ELISA plates were coated with 4 μg/ml recombinant protein (rMEPX, rMEPX_2_) in 200 μL of PBS overnight at 4 °C, which was purified from prokaryotic expression system according previous described [21]. Then blocked with 300 μL/well of 3% skimmed milk powder in PBST buffer (PBS containing 0.05% Tween 20), followed by incubation with 100 μL/well of mouse serum (1:100) at 37 °C for 1 h, and subsequently incubated with HRP-conjugated goat anti-mouse IgG (Southern Biotechnology, Birmingham, AL, USA) at a 1:2000 dilution at 37°C for 1 h. The colorimetric reaction was developed with tetramethylbenzidine (TMB) and then the absorbance at 450 nm was measured by a plate reader (Benchmark, Bio-Rad, Hayward, CA, USA) and expressed as optical density (OD) units.

### Detection of splenocyte proliferation

Splenocyte proliferation was determined by the MTT assay. Cells at a final concentration of 3×10^5^ cells/well were cultured in RPMI-1640 medium in 96-well plates in triplicate. The cultures were stimulated with ConA (positive control, final concentration 5 μg/mL) (Sigma, St Louis, MO), recombinant proteins rMEPX and rMEPX_2_ (final concentration 10 μg/mL) and BSA (negative control, final concentration 100 ng/mL) for 48 h at 37°C. Cells in culture medium only served as blank control. 20 μL (5 mg/mL) of MTT (Sigma, St Louis, MO) was added into each well and incubated for another 4 h. After 100 μL of DMSO solution was added to each well to stop the color development, plates were read at 490 nm by a microtiter plate reader (BioRad, CA, USA). Splenocyte proliferation expressed as stimulation index (SI) was calculated using the following formula: SI = (mean absorbance of each stimulation well-mean absorbance of medium)/(mean absorbance of unstimulated well-mean absorbance of medium).

### Detection of cytokines IFN-γ and IL-4 levels

The yields of IL-4 and IFN-γ in T cells were measured by intracellular cytokine staining using the double antibody sandwich method according to the manufacturer instructions of the mice IFN-γ and IL-4 ELISA kits (BOSTER Biological Technology Co. Ltd.Wuhan, China). After splenocyte cells from mice were also stimulated with different antigen, the expression levels of IFN-γ and IL-4 in the cell suspension were detected by ELISA.

### Statistical analysis

Statistical analyses were performed using GraphPad Prism version 5.0. The data was expressed as means and standard deviation. The absorbance at 450 nm of the mouse serum antibody, the cytokine IFN-γ and IL-4 levels of cell culture supernatant from the mouse spleen were analyzed using one-way ANOVA. The t-test was used to compare the mean valuesbetween groups. Differences between groups were considered significant at P < 0.05.

## Result

### Structure analysis of MEPX_2_ fragment and prediction of B cell antigen epitopes

Accessibility, variability, fragment mobility, charge distribution and hydrophilicity are important features of antigenic epitopes. The presence of flexible regions, such as coil and turn regions provides further evidence for epitope identification. In this study, the secondary structure of MEPX_2_ was predicted using the methods of Garnier and Robson [14] and Chou and Fasman [15] based on the Gn gene sequence of CCHFV YL04057 strain (Fig. 1(A)). The prediction results show that there is no transmembrane domain and signal peptide sequence in MEPX_2_. A hydrophilicity plot, flexibility plot, surface probability plot and antigenic index for the truncated protein were obtained using the methods of Kyte and Doolittle[16], Karplus and Schulz[17], Emini[18] and Jameson-Wolf[22], respectively. The analysis showed that the multi-epitope peptide is beneficial to the chimeric of the antibody, and contains the B-cell dominant epitope; the multi-epitope peptide has strong hydrophilicity and antigenicity, and the flexible region is distributed more and can be seen in the figure. A flexible region was formed at the flexible peptide added during gene construction, indicating that the multi-epitope peptide construction strategy is correct.

### Identification of recombinant plasmids pVAX-MEPX and pVAX-MEPX_2_

The recombinant plasmids pVAX-MEPX and pVAX-MEPX_2_ were digested by *Bam*HI and *Xho*I in 1% agarose gel electrophoresis. The fulllength sequence of the MEPX gene was 400 bp, the fulllength sequence of the MEPX_2_ gene was 800 bp (Fig 2(A)), consistent with the expected size. DNA sequencing confirmed that the nucleotide sequences of MEPX and MEPX_2_ in the recombinant plasmid were accurate.

**Fig 2.**
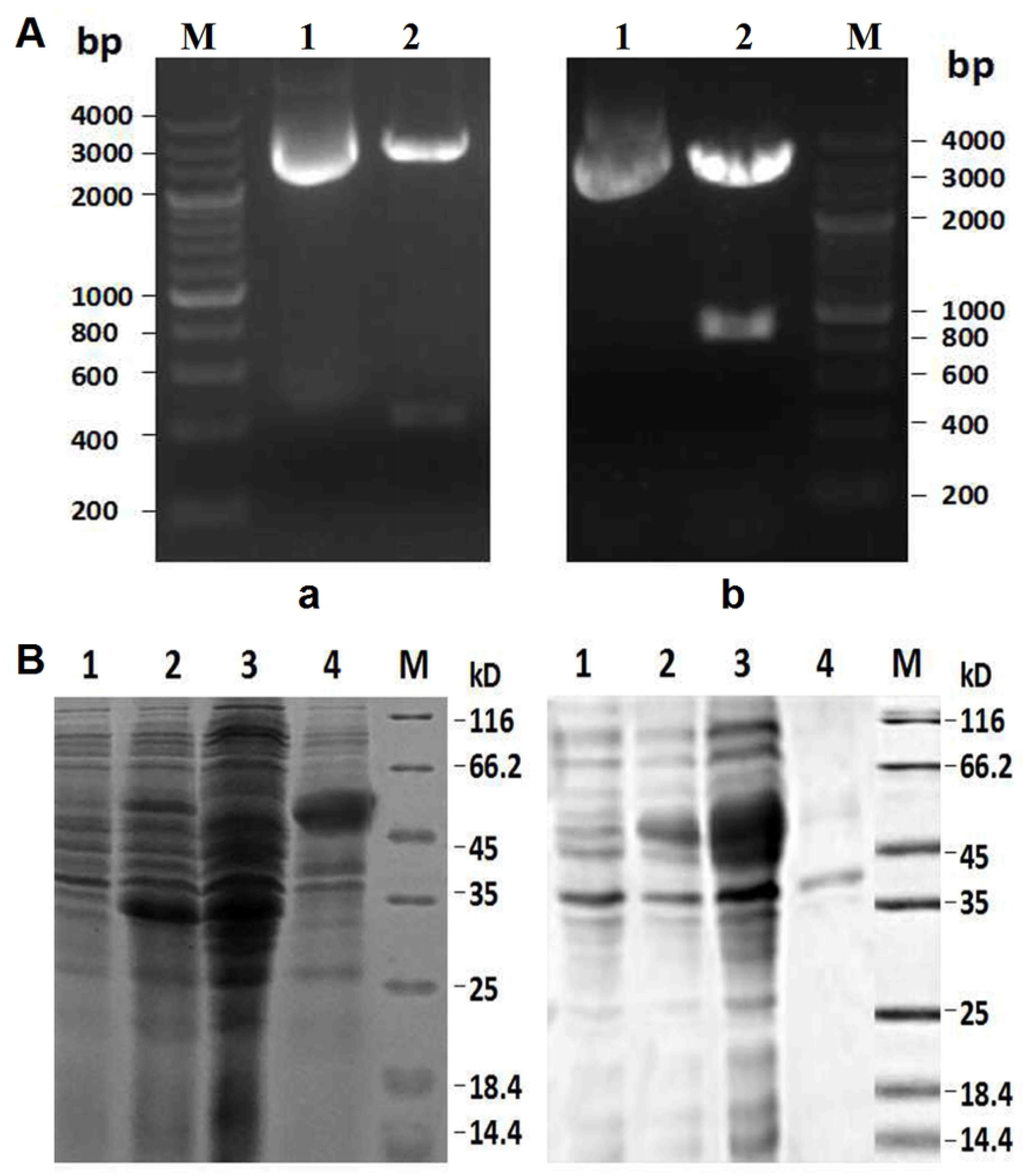
Identification of the recomibinant plasmids pVAX-MEPX and pVAX-MEPX_2_ by restriction enzyme digestion. (A) Restriction enzyme analysis of recomibinant plasmid pVAX-MEPX and pVAX-MEPX_2_, with digestion by *BamH* I and *Xho* I, follwed by 1% agarose electrophoresis. Two bands were observed for 2 recombinant plasmids, where in pVAX-MEPX is about 400 bp in size, pVAX-MEPX_2_ is about 800 bp in size. (B) SDS-PAGE analysis of rMEPX and rMEPX_2_ expression, a: rMEPX protein; b: rMEPX_2_ protein: lanes 1 and 2, lysates before and after induction; lanes 3 and 4, supernatant and inclusion bodies. M: marker.

### IFA analysis of recombinant plasmid expression in eukaryotic cells

Fluorescence microscopy revealed that pVAX-MEPX, pVAX-MEPX_2_, and pVAX-Gc-NP2 transfected cells have green fluorescence, whereas empty transfection plasmid pVAXI cells do not have any fluorescent substances (Fig 3). This result suggests that recombinant eukaryotic plasmids pVAX-MEPX, pVAX-MEPX_2_ and pVAX-Gc-NP2 (positive control) can be successfully expressed in vitro.

**Fig 3.**
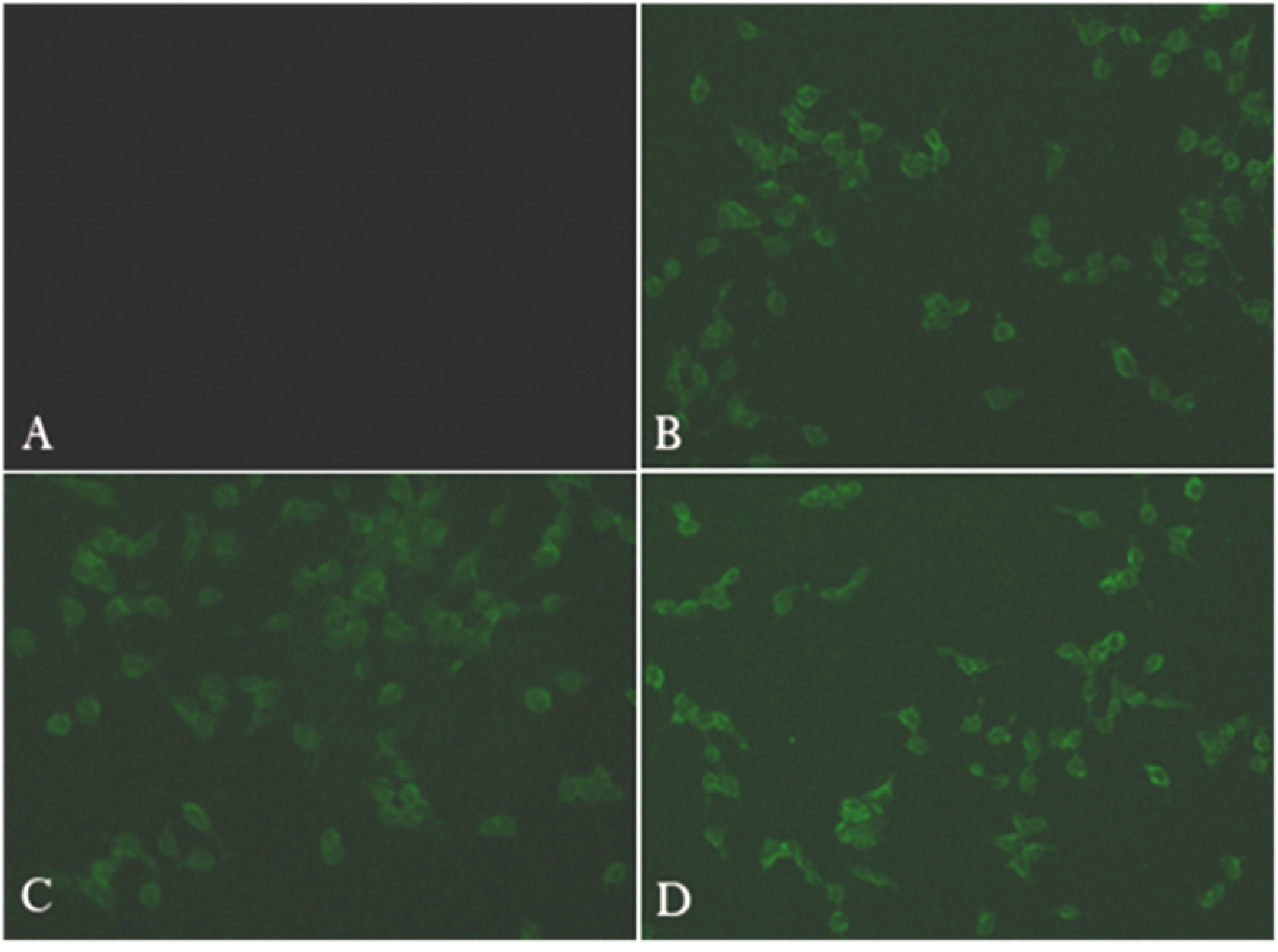
Indirect immunofluorescence identification of recombinant plasmid expression in eukaryotic 293T cells. The recombinant plasmid was transfected into 293T cells and the gene expression was analyzed by IFA analysis. (A) pVAXI, (B) pVAX-MEPX, (C) pVAX-MEPX_2_, (D) pVAX-Gc-NP2. Bars indicate 50 µm.

### Expression and purification and Western blot identification of recombinant polyepitope protein

SDS-PAGE results showed that rMEPX and rMEPX_2_ can be expressed in the supernatant (Fig 2 (B)), and can also be effectively purified using gradient elutes with imidazole (Fig 4). The protein purity can reach above 95%. Bands at approximately 34 kD and 49 kD clearly present in the Western Blot (Fig 4), indicating that the expressed rMEPX and rMEPX_2_ could specifically bind to the anti-His mouse monoclonal antibody. This result suggests that the recombinant protein rMEPX and rMEPX_2_ was expressed correctly, and recognized specifically by positive sheep serum against CCHFV antibodies, indicating that rMEPX and rMEPX_2_ have strong antigenicity.

**Fig 4.**
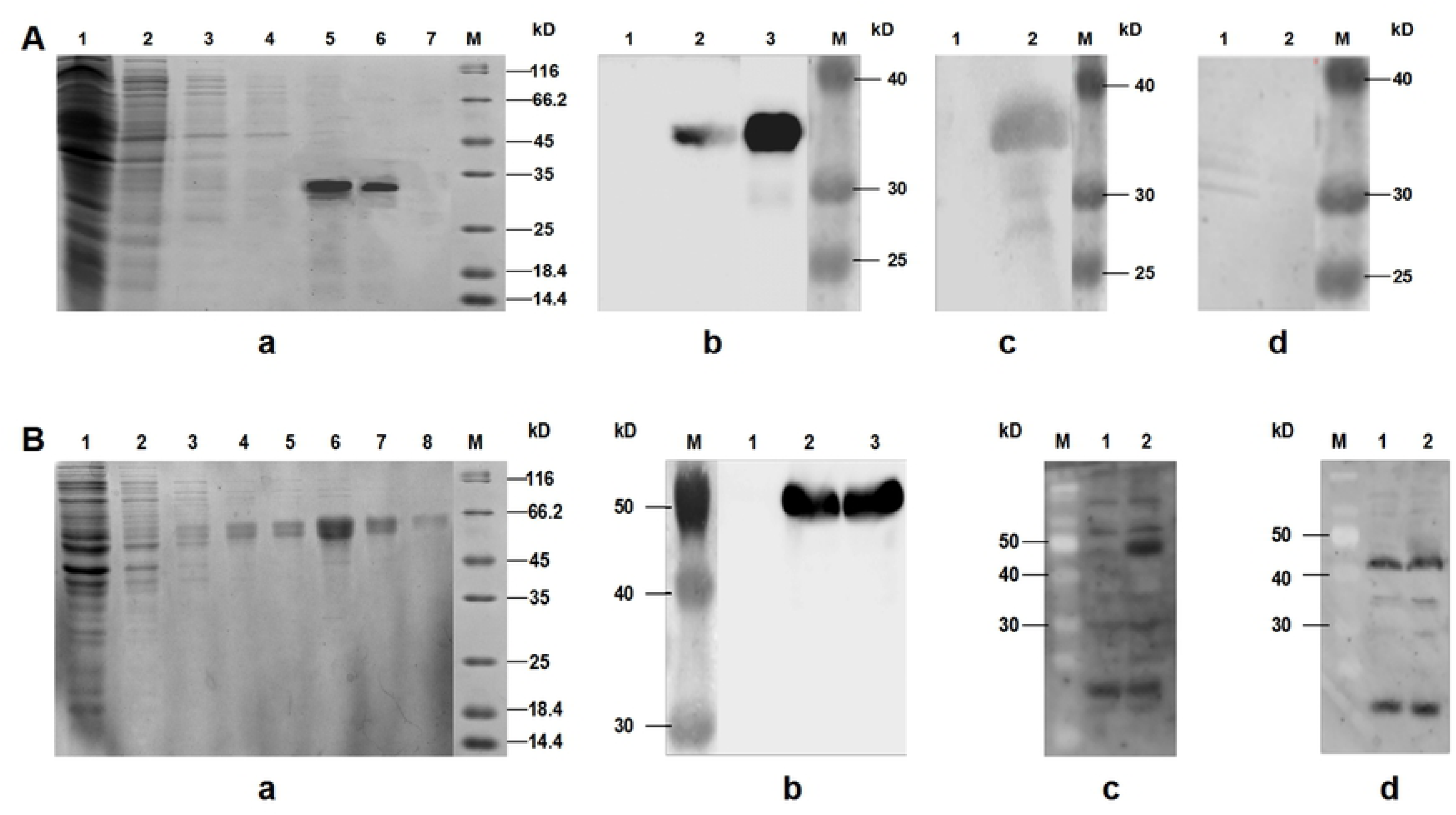
Prokaryotic expression and immunoblotting analysis of rMEPX and rMEPX2. (A) a: SDS-PAGE analysis of purification of rMEPX. 1: total protein extraction; 2-4: 25 mmol/L imidazole step-out protein sample; 5-6: 120 mmol/L imidazole eluted protein sample; 7: 500 mmol /L imidazole elution protein sample; M: standard protein marker; b: The antigenicity of rMEPX was identified by Western blotting using an anti-His mouse monoclonal antibody; c: Western blotting of rMEPX using positive sera from sheep with a confirmed history of CCHFV infection; d: Western blotting of rMEPX using sera from healthy sheep with no history of CCHFV infection used as a negative control; 7: uninduced pET-32a-MEPX/ BL21; 8: post-induction pET-32a-MEPX/BL21; 9: purified rMEPX; M: pre-stained protein Marker (B) a: SDS-PAGE analysis of purification of rMEPX_2_. 1: total protein extraction; 2-4: 25 mmol/L imidazole step-out protein sample; 5-6: 120 mmol/L imidazole eluted protein sample; 7: 500 mmol /L imidazole elution protein sample; M: standard protein marker; b: Identification of the antigenicity of rMEPX_2_ by Western blot using anti-His mouse monoclonal antibody; c: Western blotting of rMEPX_2_ using positive sera from sheep with a confirmed history of CCHFV infection; d: Western blotting of rMEPX_2_ using sera from healthy sheep with no history of CCHFV infection used as a negative control; 7: uninduced pET-32a-MEPX_2_/BL21; 8: post-induction pET-32a-MEPX_2_/BL21; 9: purified rMEPX_2_; M: pre-stained protein Marker

### Indirect ELISA detection of serum antibody levels in mice in different immunization groups

Indirect ELISA results showed that the IgG levels in the experimental groups were significantly higher than those in the control groups (Fig 5). ELISA data also showed that the experimental groups can effectively induce the humoral immune response after 14, 28, and 42 days. While the IgG levels in groups pVAX-MEPX and pVAX-MEPX_2_ were lower than those in groups rMEPX and rMEPX_2_ within 42 days, there is no significant difference in IgG levels among these groups after immunization of 42 days. The IgG level induced in group pVAX-MEPXrMEPX showed the highest level was reached at 42 days after immunization, and significantly higher than those of separate groups of DNA alone and protein alone (P < 0.01). After 42 days of immunization, the antibody titers in the mice serum immunized with pVAX-MEPX, pVAX-MEPX_2_, pVAXI, rMEPX, rMEPX_2_, PBS and co-immunization pVAX-MEPX_2_+rMEPX_2_ were 1:1600, 1:3200, 1:102400, 1:204800, 1: 204800 and 1:409600, respectively. The antibody titer of the rMEPX and rMEPX_2_ groups was 64 fold higher than that of the pVAX-MEPX and pVAX-MEPX_2_ groups.

**Fig 5.**
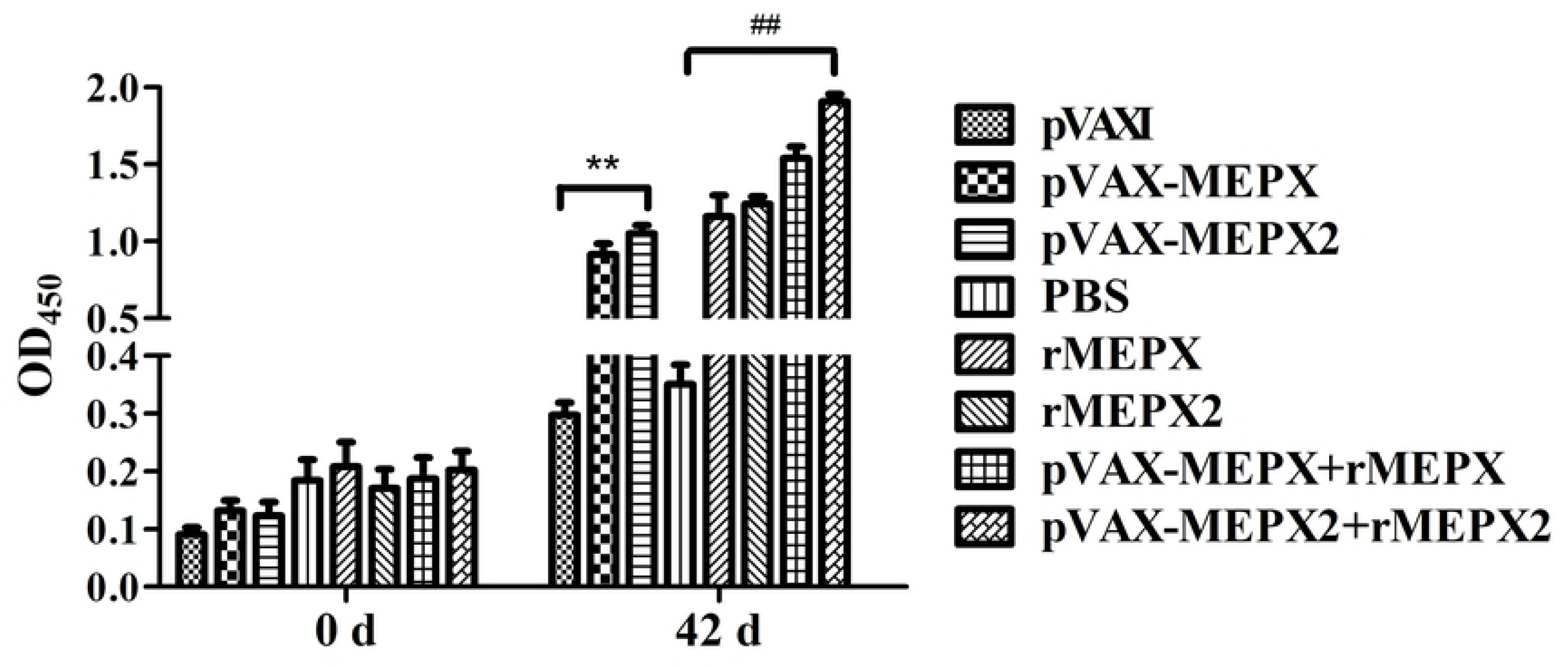
Detection special serum IgG antibodies at various times in immunized mice by ELISA. Sera was collected on day 0 and 42, and diluted (1:100) with PBST containing 5% skim milk. HRP-conjugated goat anti-mouse IgG antibody (1:2000) was used as the secondary antibody. Ab levels of IgG in sera were detected. Bar indicates the mean value of Ab levels. The PBS group and the PVAX group were used as a negative control group. **P < 0.01 vs pVAXI; ^##^P < 0.01 vs PBS.

### Lymphocyte proliferation reaction

In order to analyze the proliferation of mouse spleen T cells stimulated by each immunization group, the spleen was aseptically taken after 7 days of the third immunization. The proliferation of splenic T lymphocytes was analyzed using MTT assay. Fig. 6 exhibited that the stimulation index of DNA prime-protein vaccine boost combinatorial immunization group (pVAX-MEPX+rMEPX and pVAX-MEPX_2_+rMEPX_2_) was significantly higher than protein immunization alone, and the index is slightly higher than DNA immunization. The difference between pVAX-MEPX+rMEPX and pVAX-MEPX_2_+rMEPX_2_ group was extremely significant (P < 0.01). The stimulation index of DNA immunization group (pVAX-MEPX and pVAX-MEPX_2_ group) was significantly higher than that of protein immunization group (rMEPX and rMEPX_2_ group) (P < 0.01). In contrast, there is no significant difference in stimulation index between pVAX-MEPX group and rMEPX_2_ group.

**Fig 6.**
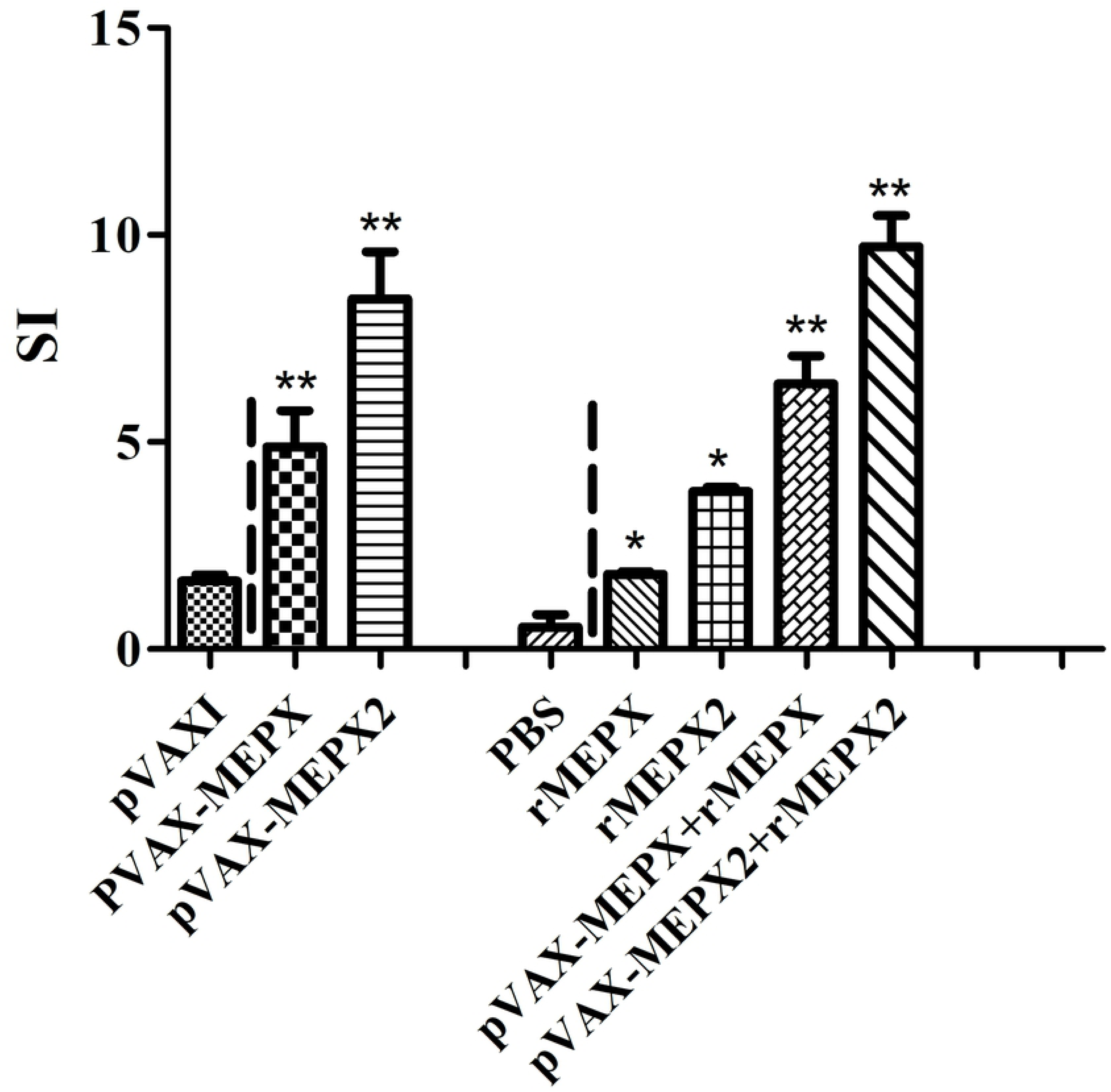
Analysis of spleen T lymphocyte proliferation in immunized mice. The immunization schedule and prime boosting of eight groups of mice are described in Materials and Methods. The immunized BALB/C mice were euthanized on day 49 and spleen cells harvested. Single cell suspensions of spleen cells (3 × 10^5^ cells/well) were stimulated in vitro with recombinant protein rMEPX and rMEPX2 (10 μg/mL) and processed for analysis of T cell proliferation as described in Material and Methods. The values are expressed as stimulation index (mean ± S.E.) obtained from the analysis of three immunized mice. **P < 0.01 vs pVAX1or PBS; *P < 0.05 vs pVAX1 or PBS.

**Fig 7.**
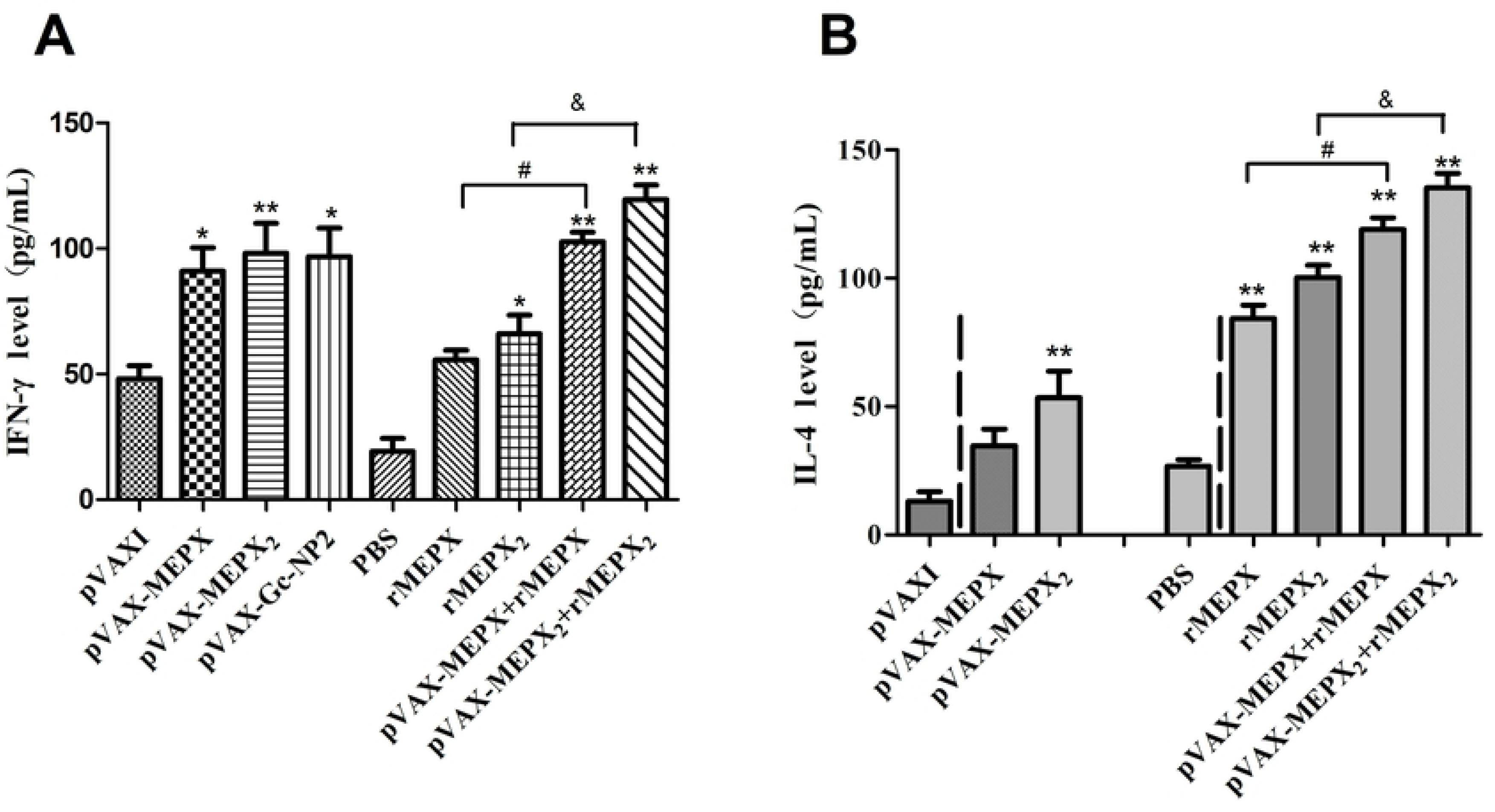
ELISA detection of IL-4 and IFN-γ expression levels in mouse spleen cell supernatant. After 7 days of the last immunization, the expression of IFN-γ and IL-4 in the supernatant of the mouse spleen T lymphocyte culture was examined. The difference between the treatment groups is statistically significant. **P < 0.01 vs pVAX1 or PBS; *P < 0.05 vs pVAX1 or PBS; #P<0.05 pVAX-MEPX+rMEPX vs rMEPX; &P<0.05 pVAX-MEPX_2_+rMEPX_2_ vs rMEPX_2∘_

### The detection of IL-4 and IFN-γ cytokine levels

After 7 days of the last immunization, the expression of IFN-γ and IL-4 in the supernatant of the mouse spleen T lymphocyte culture was examined. The difference between the treatment groups is statistically significant (P < 0.05). The IFN-γ level in the DNA alone group (pVAX-MEPX and pVAX-MEPX_2_ group) was higher than that in the protein alone group (rMEPX and rMEPX_2_ group), while the IFN-γ level in the pVAX-MEPX_2_ group combined with the rMEPX_2_ group was the highest (119.375±10.284 pg/mL), which was slightly higher than the pVAX-MEPX combined rMEPX group (102.708 ± 6.437 pg/mL).

In terms of IL-4 levels, the protein alone immunization group (rMEPX and rMEPX_2_ group) was higher than the DNA alone immunization group (pVAX-MEPX and pVAX-MEPX_2_ group), and the DNA-protein combination group (pVAX-MEPX+rMEPX and pVAX-MEPX_2_+rMEPX_2_ group) was higher than the protein alone immunization group (rMEPX and rMEPX_2_ group). The IL-4 levels in the pVAX-MEPX_2_ group combined with the rMEPX_2_ group were significantly different from those in the rMEPX and rMEPX_2_ groups (P<0.05). The highest expression IL-4 level in pVAX-MEPX_2_ combined with rMEPX_2_ group was 135.187±9.805 pg/mL, which is slightly higher than that of pVAX-MEPX combined with rMEPX group (119.035±7.768 pg/mL).

## Discussion

The method of vaccination prevention has been successfully applied to control a variety of viral diseases, such as avian influenza, Newcastle disease and Hepatitis B [23-25]. However, because CCHFV is a biological safety class IV pathogen, it requests a high level of laboratory and instrument, but suitable experimental animal models were lacked. As a result, there is currently no effective CCHF prevention, treatment, and commercially available vaccines. The multi-epitope gene-based vaccines can induce favorable and specific immune responses, and effectively avoid the side effects of other adverse epitopes in intact antigens [26]. Meanwhile, there is no risk of gene integration into host cells, largely enhancing safety [8]. Reasonable epitope design can improve immune efficacy and have the capability to focus the immune response on conserved epitopes [27]. In addition, the use of bioinformatics methods to obtain candidate epitope peptides, combined with epitope mapping to identify the fine epitopes, can facilitate the development of viral multi-epitope vaccines [12]. The use of reverse genetics technology to prepare multi-epitope gene-based vaccines provides a potential candidate tool for the CCHF prevention..

Nucleoproteins and glycoproteins are two major structural proteins of CCHFV, and play a key role in the viral infection process. Nucleoprotein NP is the main antigen that induces humoral and cellular immunity in the body [28]. The mature glycoprotein GP is very important in the process of virus infection and pathogenicity, and can induce neutralizing antibodies. Based on this, the multi-epitope vaccines design of current study includes 2 epitopes on NP2 segment, which was previously identified CCHFV YL04057 strains [19], and 4 immuno-dominant, which was highly conserved linear B-cell epitopes on glycoproteins (Gn and Gc) predicted by bioinformatics software [13]. These epitopes were used to construct the multi-epitope DNA vaccines and protein vaccines. Previous reports suggest the effect of DNA vaccine immunization alone could be limited due to the host resistance, low levels of expression, and inappropriate transfection in vivo [29, 30]. The delivery of DNA vaccines by electroporation in vivo can enhance T cell responses by increasing the efficiency of antigen-specific T cells and the ability of T cells to secrete more IFN-γ [30]. In this study, the in vivo electroporation method was used as a DNA vaccine delivery method. This study was designed in view of the DNA primed protein boost protocol and Freund’s adjuvant as a highly effective adjuvant to receive a strong immune response. Different experimental groups of DNA, recombinant protein alone, and DNA prime-protein vaccine boost combinatorial immunization were designed and performed. The results showed that the constructed multi-epitope DNA vaccines (pVAX-MEPX and pVAX-MEPX_2_) generate a higher level of IFN-γ than the control group pVAXI, indicating a significant Th1-type immune response was produced [31]. The multi-epitope protein vaccines (rMEPX and rMEPX_2_) stimulated higher levels of IL-4 compared to the PBS control group, suggesting a significant Th2-type immune response [32]. However, the DNA prime-protein vaccine boost combinatorial immunization group stimulated Th1 and Th2-type immune responses, were higher than the DNA and protein alone group, which is in good agreement with the results of Memarnejadian A et al. [33]. From the perspective of elicited antibody level, pVAX-MEPX combined with rMEPX and pVAX-MEPX_2_ combined with rMEPX_2_ immunized mice produced antibody titers up to 2.05×105 and 4.1×105, respectively. This value was much higher than recombinant immunized DNA alone, and was 2-fold higher than that of recombinant protein alone group. This finding confirms that the DNA prime-protein boost strategy can enhance antibody levels, which is consistent with Liang’s study result [34]. The mechanism may be enhanced by the stimulation of antigen-specific memory B cells, followed by the protein immune coordination through the process of DNA primary immunization [35]. Meanwhile, the prime-boost immunization strategy amplifies selective B or T cells with greater affinity, increasing the number of memory cells specific for common antigens [36]. We found that the levels of both cellular and humoral immune responses produced by mice immunized with rMEPX_2_ and pVAX-MEPX_2_ combined with double-copy multi-epitope recombinant DNA (pVAX-MEPX_2_) and pVAX-MEPX_2_ were significantly higher than those of pVAX-MEPX, rMEPX and pVAX-MEPX combined with rMEPX. This result also revealed that high epitope densities can induce the production of high-affinity antibodies. Yan Xuhua et al. [37] found that the increase in the number of epitope repeats can enhance the recognition and interaction of B-cell surface antibody with multi-epitope peptides. Our previous study also confirmed that the double-copy multi-epitope peptide has better antigenicity [13]. However, since CCHFV is a highly pathogenic virus and requires a high level of safety laboratory, there is no cell model and animal model to detect CCHFV in China. Therefore, this study did not further detect whether the antibodies induced by immunized mice have neutralization and protection effects.

In conclusion, our results revealed that the multi-epitope DNA vaccine and recombinant multi-epitope protein vaccine could stimulate mice to produce specific humoral and cellular immune responses. Double-copy multi-epitope vaccines generated better cellular and humoral immune responses by DNA prime-protein vaccine boost combinatorial immunization strategy. The pVAX-MEPX_2_ gene prime followed by boosting recombinant protein rMEPX_2_ immunization induced the strongest antigen-specific immune effect, would be expected to serve as vaccine candidates for the prevention and control of CCHFV. Further investigation by establishing the CCHFV animal model is needed to clarify whether the antibodies have neutralizing activity and neutralization protection.

## Acknowledgments

This work was supported by the grants from the National Science Foundation of China (No. 81460303, 81760365), the Ministry of Science and Technology of China (No. 2013FY113500), and the Science Research Key Project of Xinjiang Education Department (No.XJEDU2019I06).

